# SRSF12 is a primate-specific splicing factor that induces a tissue-specific gene expression program

**DOI:** 10.1101/2025.07.25.666902

**Authors:** Jimmy Ly, Sarah L. Cady, Ekaterina Khalizeva, Sofia Haug, Iain M. Cheeseman

## Abstract

Alternative splicing expands proteomic diversity and is tightly regulated by splicing factors, including the serine/arginine-rich (SR) protein family. Here, we analyze the poorly characterized protein SRSF12. Although SRSF12 is conserved across vertebrates, it is poorly expressed in most mammals, and we find that SRSF12 knockout mice do not display overt physiological or transcriptomic alterations. In contrast, SRSF12 is more highly expressed in primates where it is predominantly transcribed in the testes, oocytes, and brain. SRSF12 co-localizes with other splicing components to nuclear speckles and interacts with core splicing factors in cultured human cells. Strikingly, ectopic expression of SRSF12 in human cells induces widespread transcriptional changes, activating meiosis-, testis- and brain-specific genes. SRSF12 overexpression also leads to mitotic arrest and cell death, phenotypes that require both its structured RNA recognition motif and intrinsically disordered arginine/serine-rich C-terminal domain. Together, our results suggest that SRSF12 has evolved primate-specific expression to regulate testis- and brain-specific genes.

## Introduction

Alternative splicing is a critical step in the regulation of gene expression that acts to increase proteome complexity and modulate regulatory regions of mRNAs (Kjer-Hansen and Weatheritt, 2023; Kornblihtt et al., 2013; Marasco and Kornblihtt, 2023; Wright et al., 2022). Alternative splicing is also differentially controlled across a wide range of conditions, with a substantial change in the splicing landscape across tissues (Garcia-Perez et al., 2023; Naro et al., 2021; Yeo et al., 2004; Zhang et al., 2016). The core splicing machinery, known as the spliceosome, consists of hundreds of protein components and 5 non-coding small nuclear RNAs (Shi, 2017; Wilkinson et al., 2020). One major class of splicing regulators that functions in both constitutive and alternative splicing is the serine/arginine-rich (SR) RNA-binding protein family, which consists of 12 proteins in humans (SRSF1 – SRSF12) (Jeong, 2017; Zheng et al., 2020). SR proteins generally enhance splicing when bound to exons and inhibit splicing when bound to introns (Erkelenz et al., 2013). In addition to their role of SR proteins in alternative splicing, this family of proteins also has established roles in regulating transcription (Li et al., 2023). Therefore, alterations to the expression of SR proteins can influence both transcription and the alternative splicing program. Some SR proteins may also be required for cell proliferation. For example, knockdown of SR proteins has been shown to disrupt cell cycle progression, including the induction of mitotic errors (de Wolf et al., 2021; Funk et al., 2022; Pellacani et al., 2018), whereas overexpression of SR proteins is correlated with several diseases (Anczukow et al., 2012; Khalife et al., 2025; Li et al., 2023; More and Kumar, 2020).

In our search for splicing regulators that could contribute to tissue-specific patterns of alternate mRNA splicing (Ly et al., 2024a), we identified SRSF12 as displaying differential expression across tissues. Although the function of most SR proteins has been well documented, the function of SRSF12 remains understudied. Based on studies using in vitro extracts, prior work suggested that SRSF12 act as a repressor of splicing by antagonizing the function of SRSF proteins (Cowper et al., 2001). Similarly, a recent report demonstrated that expression of SRSF12 causes skipping of the alternative exon 4 of CLK1, highlighting an interesting autoregulatory loop (Shkreta et al., 2024). Some reports suggested possible roles for SRSF12 in tumorigenesis (Yang et al., 2025) or the immune response (Wilton et al., 2023; Zhang et al., 2024). However, the function of SRSF12 in both alternative splicing or transcription has not been well studied in a cellular or organismal context. Here, we show that SRSF12 is uniquely expressed in the testis and brain of primates. Although we do not observe strong phenotypes for SRSF12 knockouts in murine models, we found that ectopic SRSF12 expression in human HeLa cells uniquely induces the expression of tissue-specific genes. SRSF12 overexpression results in dramatic cell cycle defects and arrests cells in mitosis. Through a structure-function analysis, we show that both the structured N-terminal RNA recognition motif and unstructured C-terminal SR-rich region are required for its activity when overexpressed in HeLa cells. Overall, our work implicates SRSF12 in controlling tissue-specific gene expression programs and highlights the evolution of its cell-type specificity.

## Results

### SRSF12 is a brain and germline specific splicing factor in primates

To gain insight into the regulation of SRSF12, we analyzed its expression patterns across tissues and evolutionary lineages. The SRSF12 gene is conserved across vertebrates—including primates, rodents, amphibians, and fish—but is absent from invertebrate models such as *Drosophila melanogaster* and *Caenorhabditis elegans* (Fig. 1A). Between humans and mice, the protein sequence is highly conserved (93% amino acid sequence identity), with the most conserved region being the RNA-recognition motif at the N-terminus (Fig. S1A).

**Figure 1.**
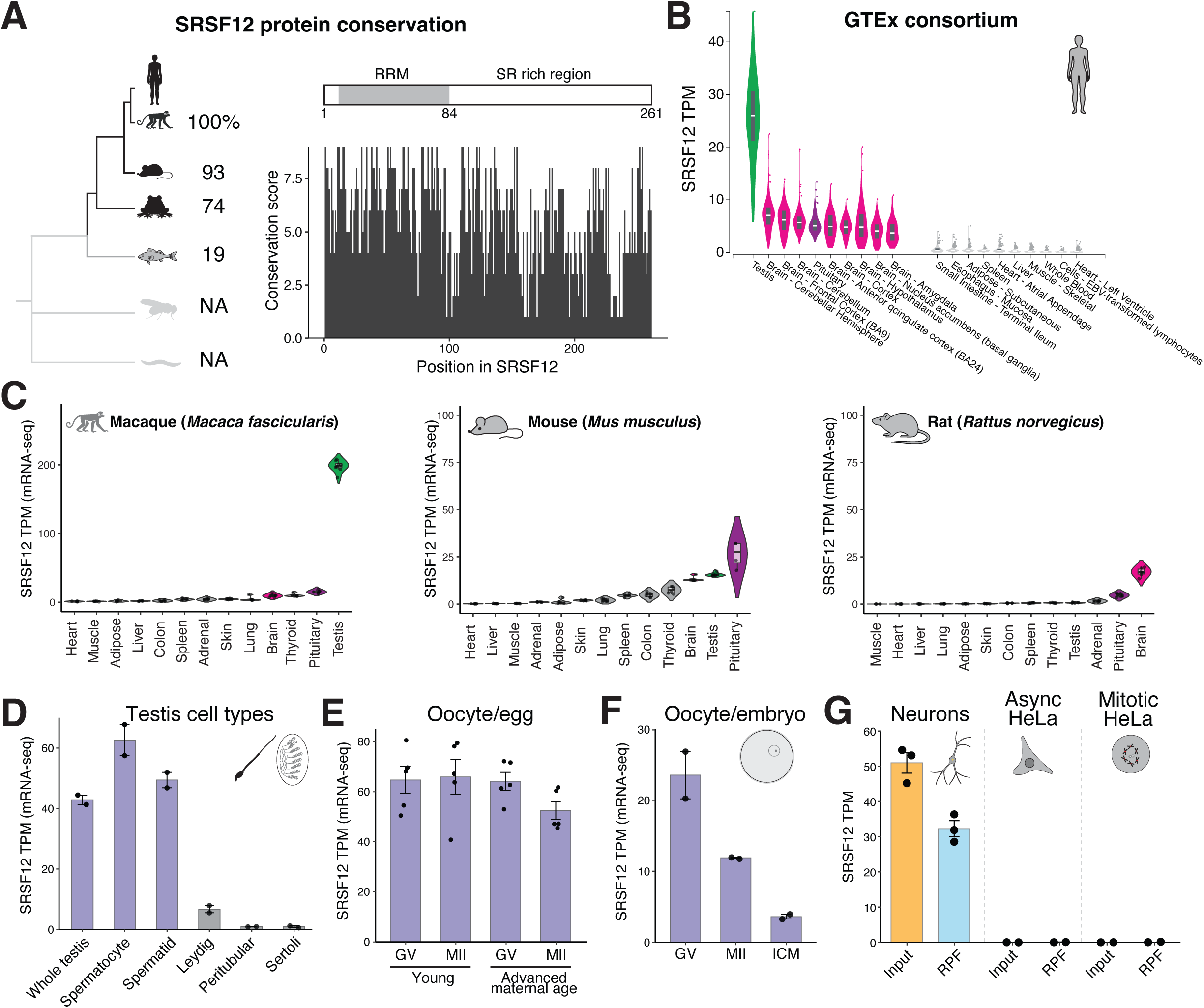
SRSF12 is a meiosis and brain specific factor in primates. (**A**) Conservation of SRSF12 in metazoan model organisms along with SRSF12 sequence identity (left). SRSF12 conservation scores per position from ConSurf (Yariv et al., 2023). (**B**) SRSF12 TPM from indicated human tissues from the GTEx consortium (Consortium, 2013). Shown are the top 10 and bottom 10 SRSF12 expressing tissues. (**C**) SRSF12 TPM from indicated tissues from various mammals (Naqvi et al., 2019). (**D**) SRSF12 TPM from specific cell types from human testis (Jegou et al., 2017). (**E**) SRSF12 TPM from human GV oocytes and meiosis II eggs from young and advanced maternal age donors (Reyes et al., 2017). (**F**) SRSF12 TPM from human GV oocytes and early embryos (Wu et al., 2018). (**G**) SRSF12 TPM from input RNA-seq and ribosome protected fragments (RPF) in ribosome profiling experiments from NGN2-derived neurons (Duffy et al., 2022), interphase and mitotic HeLa cells (Ly et al., 2024b). Bars represent mean and error bars indicate standard error of the mean.

In humans and monkeys, SRSF12 displays striking tissue specificity, with high expression in the testes of both humans (TPM = 26) (Consortium, 2013) and macaques (TPM = 197) (Naqvi et al., 2019), moderate expression in the brain (TPM ∼ 10 in human and macaques), and negligible expression in other tissues (Fig. 1B-C). Within the human testis, SRSF12 is highly expressed in meiotic spermatocytes (TPM = 57.4) and mature spermatids (TPM = 97.4) but not the somatic testes cells such as Leydig (TPM = 5.5), Sertoli (TPM = 0.6), or Peritubular cells (TPM = 0.9; Fig. 1D) (Jegou et al., 2017). Consistent with its expression during meiosis, SRSF12 is present in human germinal vesicle–arrested oocytes and metaphase II eggs, with expression levels remaining stable with age (Fig. 1E) (Reyes et al., 2017). In contrast, human SRSF12 mRNA expression declines >6-fold during early embryonic development from oocytes to embryonic day 5 blastocysts (Fig. 1F) (Wu et al., 2018). To assess whether SRSF12 is also expressed in neurons, we analyzed RNA-seq and ribosome profiling data from NGN2-induced neurons derived from human iPSCs (Duffy et al., 2022). These data revealed that SRSF12 is both transcribed and actively translated in neurons but not in cancer cell models such as HeLa cells (Fig. 1G) (Ly et al., 2024b). Thus, SRSF12 is preferentially expressed in primate brains and testis.

Despite strong sequence conservation between human and mouse SRSF12 (Fig. 1A), rodents show minimal expression of SRSF12 across all tissues and cell types examined, with very modest expression in the brain, pituitary, and testis (Fig. 1C) (Naqvi et al., 2019). Together, these findings suggest that SRSF12 is broadly conserved among vertebrates, but that its robust and brain- and testis-specific expression likely represent a relatively recent evolutionary adaptation in primates.

### SRSF12 is dispensable in mouse models

Although SRSF12 expression is extremely low in rodents (Fig. 1C), given the potential for tissue-specific contributions and the challenges of genetic studies in primates, we next assessed its function in mouse models. We generated a SRSF12 knockout allele using frameshift mutations in the N-terminal RNA recognition motif (Fig. S1B-C). Homozygous SRSF12 knockout mice developed normally, were fertile, and showed no overt phenotypes (Fig. S1D-G). Transcriptomic analysis of brain and testis revealed only 37 significant differences between knockout and wild-type animals across these two tissues (Fig. S1H-I; Table S1). Similarly, differential exon usage analysis using DEXSeq (Anders et al., 2012) revealed only 6 and 2 significantly differentially used exons in testis and brain, respectively (Table S1). These transcriptomic results are consistent with SRSF12 being dispensable for mouse development and viability. Given the distinct expression profiles that we observed between primates and rodents, the absence of a mouse knockout phenotype suggests that the functional contributions of SRSF12 may be limited to primates. Alternatively, there may be an SRSF12 paralog or alternative pathway that can compensate for loss of SRSF12 function in mice.

### SRSF12 is hyperphosphorylated and interacts with the core splicing machinery along with transcription factors in cell culture

We next sought to characterize the function of SRSF12. To do this, we expressed SRSF12 tagged with a C-terminal GFP under a dox-inducible promoter in HeLa cells and performed immunoprecipitations and mass spectrometry. Through this analysis, we found that SRSF12 is highly phosphorylated at its unstructured C-terminal serine and arginine-rich tail (Fig. S2A), possibly by the kinase, CLK1 (Shkreta et al., 2024). Furthermore, SRSF12 strongly interacted with several components of the splicing machinery such as SRSF1, other SRSF and hnRNP family proteins, the m6A reader YTHDC1 (Xiao et al., 2016), and other RNA-processing factors (Fig. 2A-B; Fig. S2B). SRSF12 also interacted with several transcriptional regulators, such as ZNF768 (Rohrmoser et al., 2019) and ZC3H4 (Estell et al., 2021).

**Figure 2.**
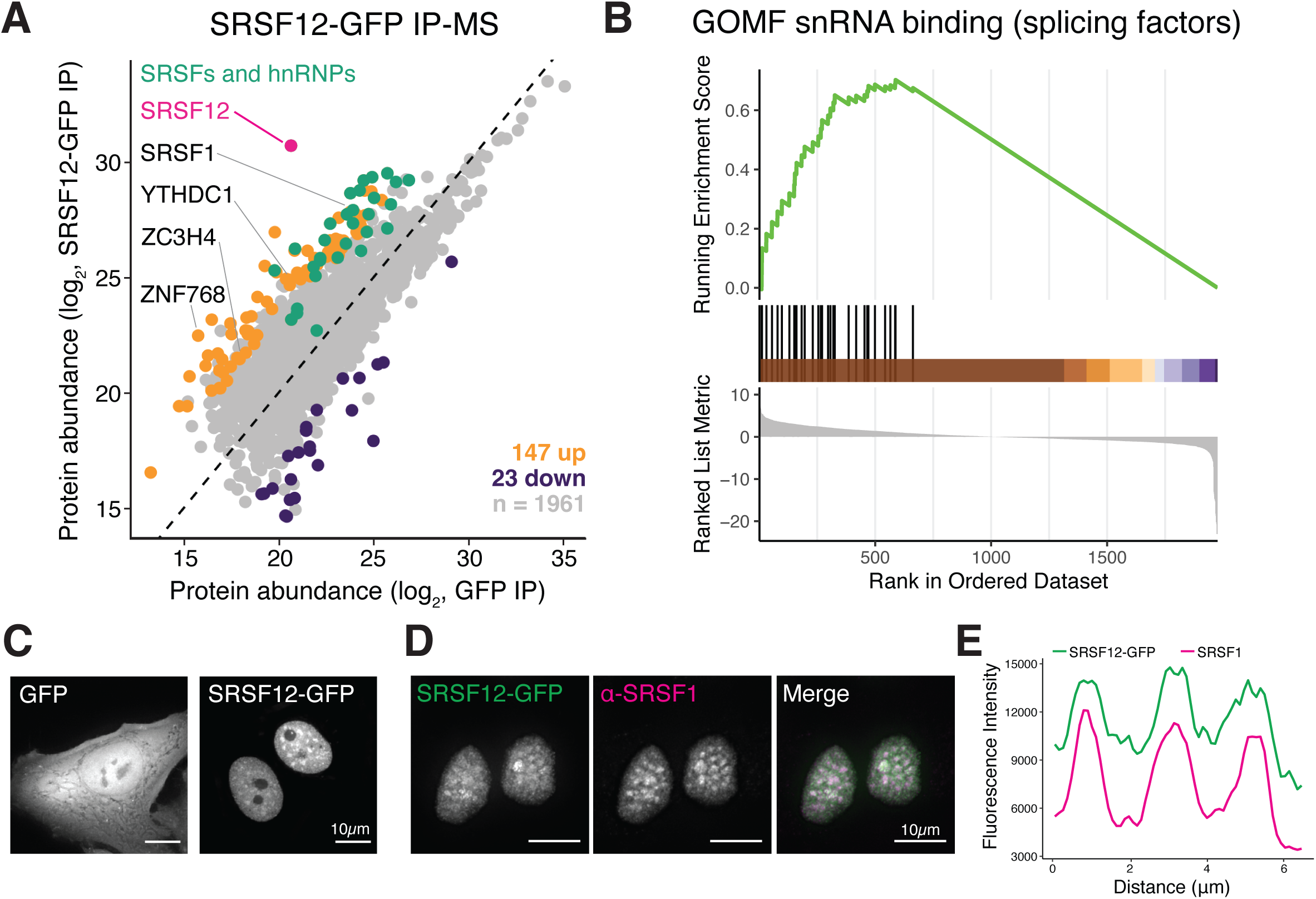
SRSF12 interacts with core splicing components. (**A**) Scatterplots showing the relative abundances of the proteins detected in GFP (x-axis) and SRSF12-GFP (y-axis) IP-MS. Proteins significantly enriched in SRSF12-GFP immunoprecipitates are shown in orange, whereas significantly depleted proteins are shown in purple. Abundances represent the average of 2 biological replicates. Statistical significance is defined as >4-fold change and p-value ≤ 0.05. (**B**) Gene set enrichment analysis plot from SRSF12-GFP IP-MS experiments, highlighting the enrichment of splicing factors. (**C**) Live-cell imaging of inducibly expressed GFP and SRSF12-GFP in HeLa cells. (**D**) Immunofluorescence of SRSF12-GFP expressed in HeLa cells, co-stained with SRSF1 antibody. (**E**) Line plot highlighting the co-localization of SRSF12-GFP and SRSF1 in nuclear foci.

To confirm the interactions with the core splicing machinery, we tested SRSF12-GFP localization. We found that SRSF12-GFP localized to the nucleus and formed prominent foci (Fig. 2C). Co-immunofluorescence with canonical splicing factors revealed that these foci overlap with classical nuclear speckles marked by SRSF1, consistent with the SRSF12 interaction profiling (Fig. 2D-E). Overall, these data suggest that, when expressed in HeLa cells, SRSF12 is a highly phosphorylated protein that co-localizes and interacts with the core splicing machinery.

### SRSF12 overexpression in somatic cells induce cell death through mitotic arrest

We next evaluated how ectopic expression of SRSF12 affects cellular fitness in HeLa cells using pooled competitive growth assays. For these experiments, we combined doxycycline-inducible SRSF12-GFP cells were equal ratios of inducible mCherry-expressing control cells and tracked the relative ratio of GFP-positive and mCherry-positive cells over time. SRSF12 expression led to a sharp decline in fitness, indicating that SRSF12 expression is highly toxic in HeLa cells (Fig. 3A). To understand the molecular mechanism underlying this death, we first analyzed cell cycle progression. Ectopic expression of SRSF12-GFP induced an accumulation of cells in the G2/M phase of the cell cycle (Fig. 3B). To distinguish G2 from a mitotic arrest, we performed flow cytometry using the mitosis-specific marker, phosphorylated histone H3 at serine 10. We found that SRSF12 expression led to an accumulation of cells in mitosis (Fig. 3C). Microscopy-based analysis of DNA morphology revealed that SRSF12-overexpressing cells exhibited a high frequency of metaphase chromosome misalignment defects, along with an increased incidence of multinucleated cells (Fig. 3D-E). We also observed similar results upon expression of the mouse SRSF12 protein sequence in HeLa cells (Fig. S3). In addition, this phenotype was independent of the transgene expression system (Fig. S3). Together, these findings suggest that ectopic expression of mammalian SRSF12 in proliferating somatic cells is toxic and disrupts proper mitotic progression.

**Figure 3.**
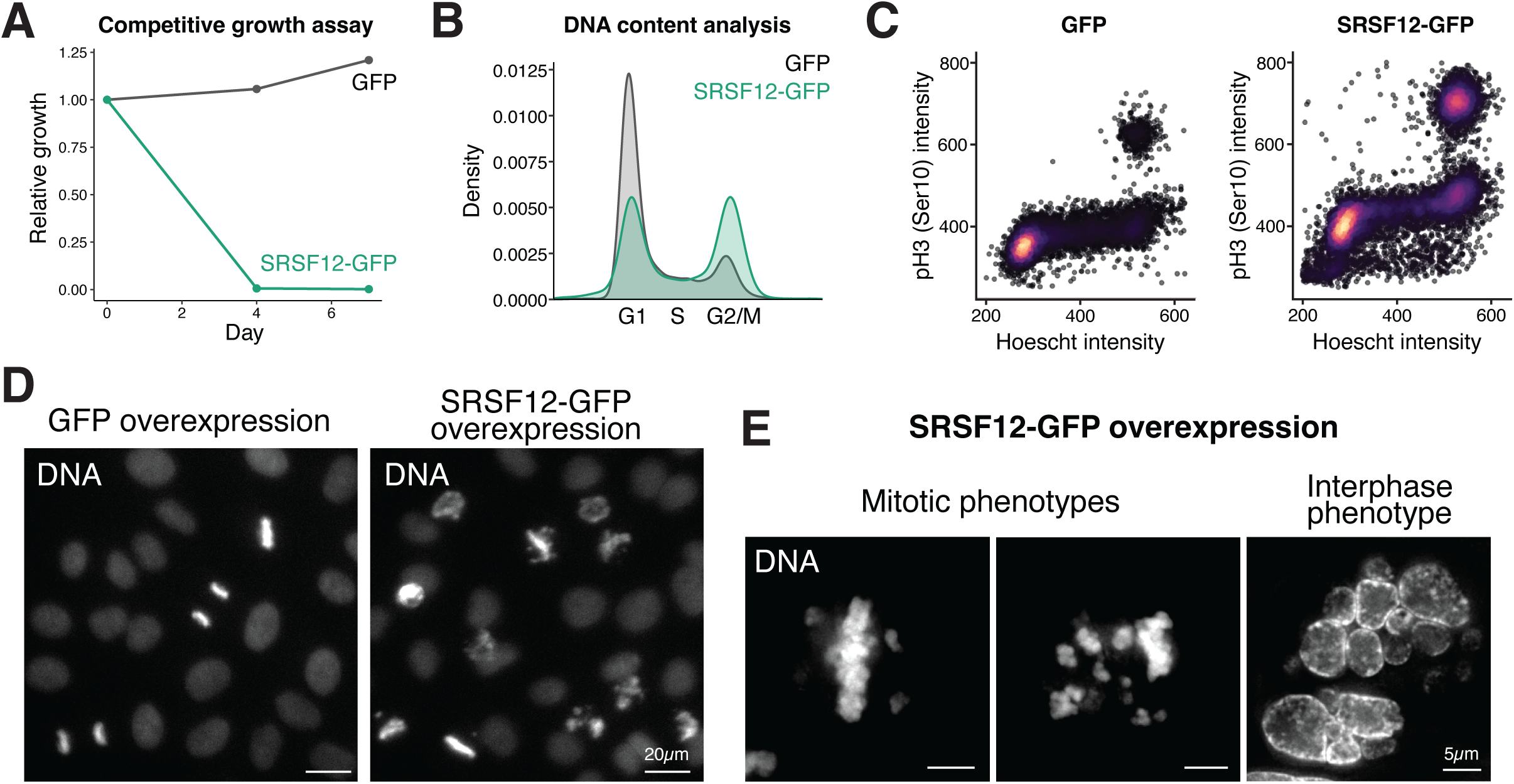
Overexpressing SRSF12 in proliferating somatic cells induces cell death through mitotic arrest. (**A**) Competitive growth assay comparing cells expressing SRSF12-GFP or GFP to mCherry-expressing reference cells. (**B**) DNA-content analysis of HeLa cells expressing SRSF12-GFP or GFP. (**C**) Scatterplot showing DNA content (x-axis) and phosphorylated histone H3 serine 10 (y-axis) in HeLa cells expressing SRSF12-GFP or GFP. (**D**) Low magnification live cell imaging DNA in control GFP and SRSF12-GFP expressing HeLa cells, highlighting the increased proportion of mitotic cells. (**E**) High magnification live-cell imaging of DNA in SRSF12-GFP expressing HeLa cells.

### Both the structured and unstructured regions of SRSF12 are required for its activity

SRSF12 contains an N-terminal RNA recognition motif (RRM, 1-84) and an intrinsically disordered C-terminal domain rich in serine and arginine residues (85-261, Fig. 4A). To determine whether both domains are necessary for the phenotypes observed upon SRSF12 expression in HeLa cells, we tested the effects of expressing either the N- or C-terminal domain alone, each fused to a C-terminal GFP tag. Both truncated proteins localized to the nucleus and formed the characteristic nuclear foci observed for full-length SRSF12 (Fig. S5A). However, in contrast to the full-length protein, expression of either domain alone did not lead to a mitotic arrest (Fig. 4B). These results indicate that either the N- or the C-terminal domains is sufficient for SRSF12’s nuclear compartmentalization but that both domains required for its activity in inducing a mitotic arrest.

**Figure 4.**
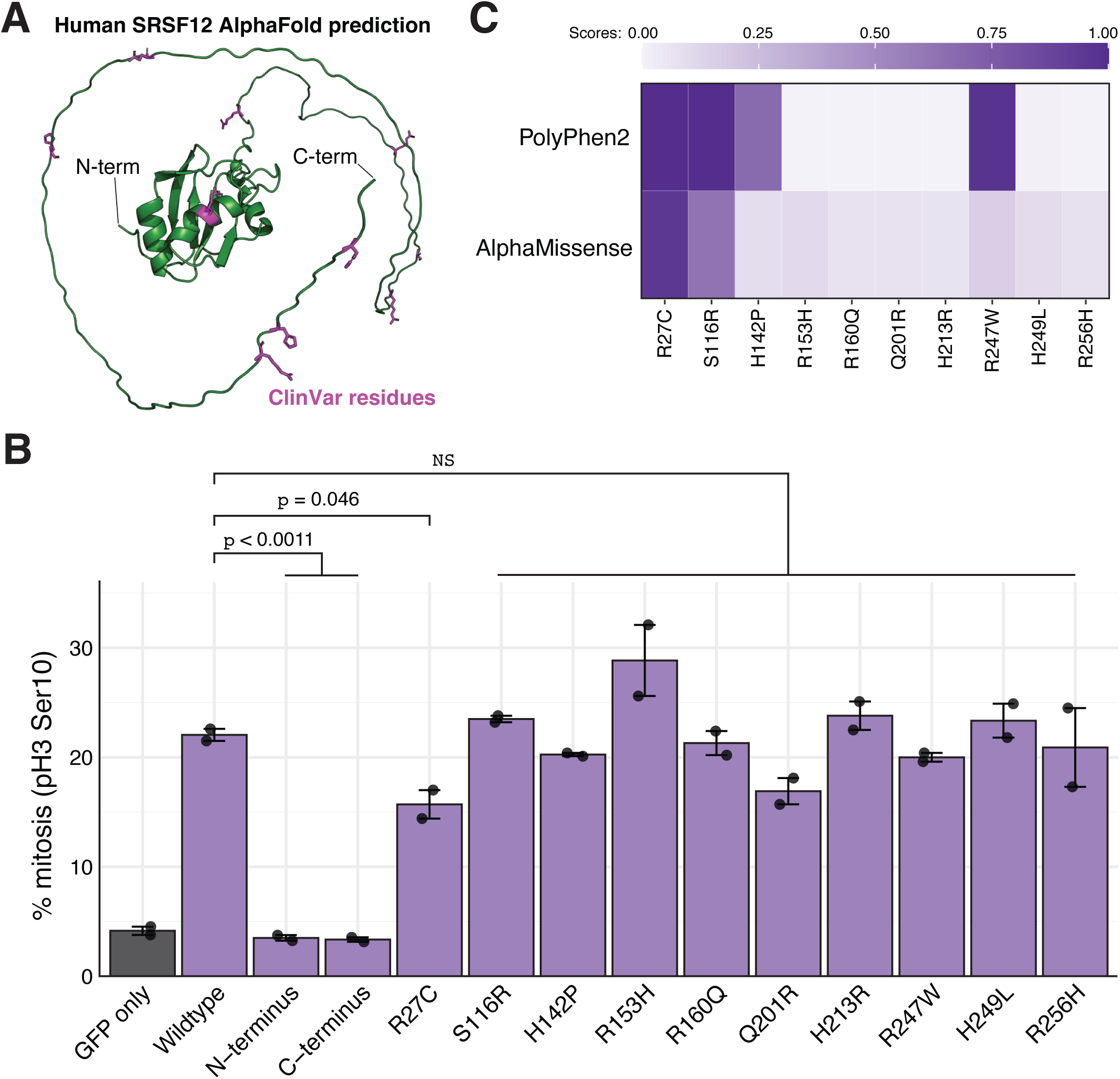
Both the structured and unstructured regions of SRSF12 are required for function. (**A**) Alphafold structure of human SRSF12. Magenta represents residues that are mutated in ClinVar (Landrum et al., 2018). (**B**) Bar graph showing the % of cells in mitosis as measured by histone H3 serine 10 phosphorylation flow cytometry in the indicated cell lines. N = 2 biological replicates. Student’s T-tests were used for statistical tests. Error bars indicate standard error of the mean. (**C**) Heatmap showing the PolyPhen2 and AlphaMissense scores for the indicated ClinVar mutations.

Given the potential link between SRSF12 and disease (Yang et al., 2025; Zhang et al., 2024), we also modeled all 10 missense variants annotated in the ClinVar database (Landrum et al., 2018). Although these variants were found in patients, no associated disease phenotypes were reported, leaving SRSF12’s role in disease unclear. To assess their functional impact, we expressed each variant in HeLa cells and tested their ability to induce a mitotic arrest (Fig. 4B-C; Fig. S4B-C). Most variants had little effect, suggesting that they do not disrupt SRSF12’s core activity in this context. However, the R27C variant caused showed a modest but significant reduction in mitotic arrest frequency, possibly implicating residue R27 in SRSF12 function (Fig. 4B). Consistently, R27C received the highest pathogenicity scores from AlphaMissense and PolyPhen2 (Fig. 4C). Based on ectopic expression assays in HeLa cells, these ClinVar mutations do not appear to represent loss-of-function or damaging alleles.

### SRSF12 expression in HeLa cells modulates splicing and 3’ end processing

Finally, to understand the molecular basis of this toxicity and mitotic arrest upon SRSF12 ectopic expression, we performed RNA-seq of SRSF12-expressing HeLa cells and analyzed RNA processing. Based on the contributions of other SR proteins to alternative splicing (Jeong, 2017; Zheng et al., 2020), we first analyzed differential exon usage upon SRSF12 expression in HeLa cells. Using DEXSeq (Anders et al., 2012), we found a total of 162 differentially used exons in HeLa cells expressing SRSF12 (Fig. 5A; Table S2). For example, SRSF12 expression enhanced the inclusion of MAP4 exon 9 (ENST00000683076, chr3: 47,912,421-47,909,038; Fig. 5B), resulting in the addition of 1129 amino acids. We also observed increased retention of ATP5IF1 intron 2 (Fig. 5C) upon SRSF12 expression, suggesting that SRSF12 can regulate both exon inclusion and intron retention. Additionally, for a subset of genes, SRSF12 expression led to a pronounced depletion of reads at the 3′ ends of mRNAs, including MECP2 (Fig. 5D) and CEP78 (Fig. 5E), among others, implying a potential role for SRSF12 in 3′ end processing. Thus, in addition to the role of SRSF12 in repressing exon inclusion (Cowper et al., 2001; Shkreta et al., 2024), our results suggest that SRSF12 expression in HeLa cells may have a more complex role in mRNA processing including exon inclusion and 3’ end formation, highlighting a multifaceted role for this protein.

**Figure 5.**
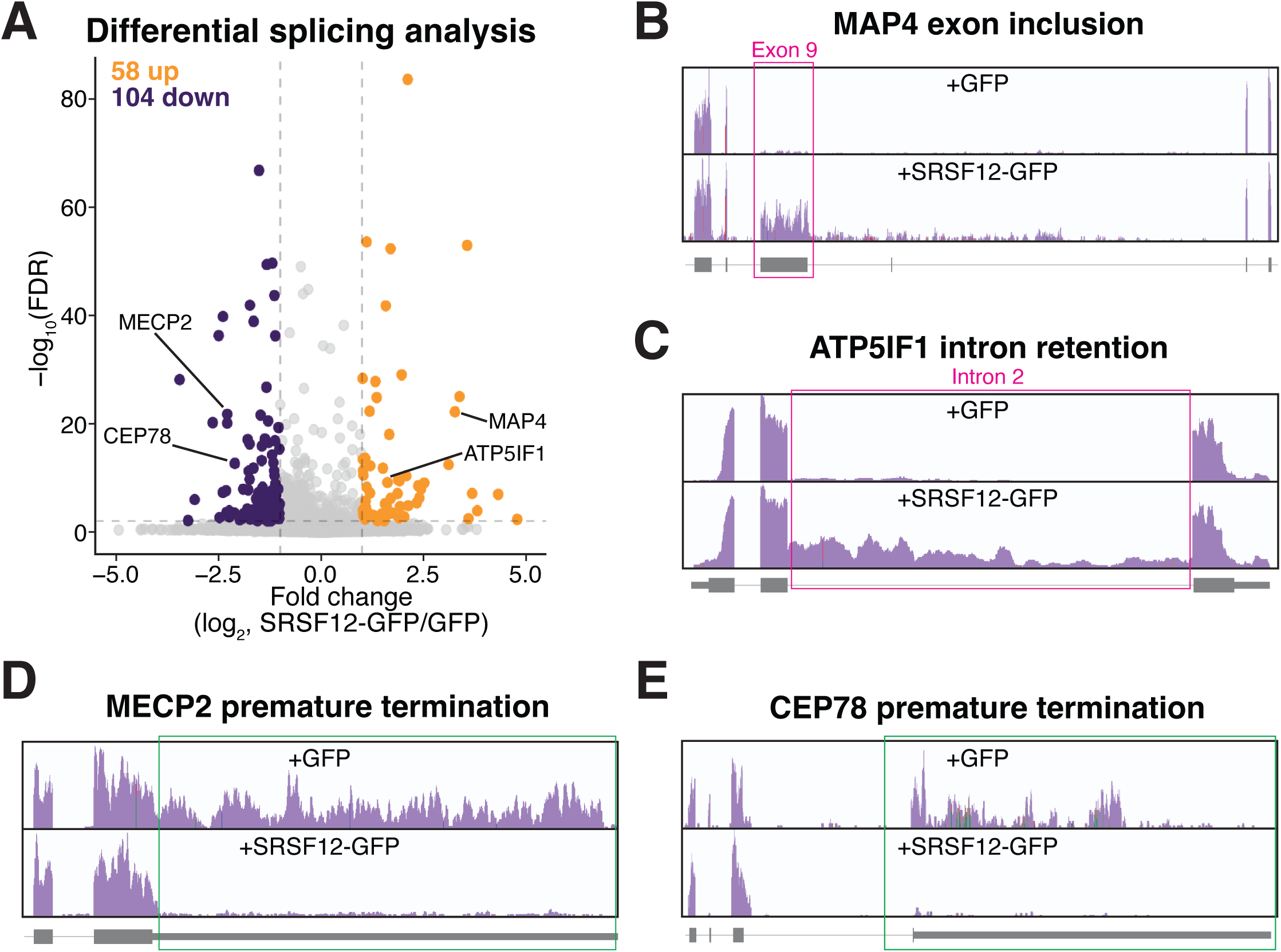
SRSF12 expression modulates splicing and 3’ end processing. (**A**) Volcano plot showing differential exon analysis in cells expressing SRSF12-GFP compared to GFP using DEXSeq. Orange, significantly upregulated exons in SRSF12-GFP; purple, significantly downregulated exons in SRSF12-GFP expressing cells. Exons with false discovery rate (FDR) < 0.01 and a fold change (FC) > 2 were called as significant. n = 3 biological replicates. (**B-E**) RNA-sequencing coverage plots for the indicated genes. The magenta or green box indicates the differentially processed region in SRSF12-GFP expressing cells.

### Ectopic expression of SRSF12 induces the expression of tissue-specific genes

Our work demonstrates a role for SRSF12 in regulating splicing and transcription termination. We next analyzed the consequences of SRSF12 on mRNA levels. Remarkably, SRSF12 expression induced a widespread change in gene expression with 412 genes downregulated and 282 upregulated (Fig. 5A-B). Notably, genes upregulated by SRSF12 overexpression tended to be lowly expressed in HeLa cells (Fig. 5B; Fig. S5A) and enriched for tissue-specific genes (Fig. S5B-C). These tissue-specific genes include meiotic-enriched transcripts such meiosis specific aurora kinase (AURKC, (Nguyen et al., 2018; Yang et al., 2010)) and the meiosis-specific telomere-associated protein (TERB1, (Shibuya et al., 2014)), testis specific proteins such as Sperm Acrosome Developmental Regulator (SPACDR/C7orf61 (Wu et al., 2024); Fig. 6C-D), and brain-specific genes such as Proline rich 18 (PRR18; Fig. 6E-F). SRSF12 expression also induced the expression of Collagen Type XX Alpha 1 Chain (COL20A1), a gene that has enriched expression in both the testis and brain based on the GTEx consortium (Consortium, 2013) (Fig. S5D). The selective induction of brain and testis gene programs following SRSF12 induction aligns with its primate-specific expression in these tissues, which may suggest a role of SRSF12 in regulating tissue-specific transcription (Fig. 1).

**Figure 6.**
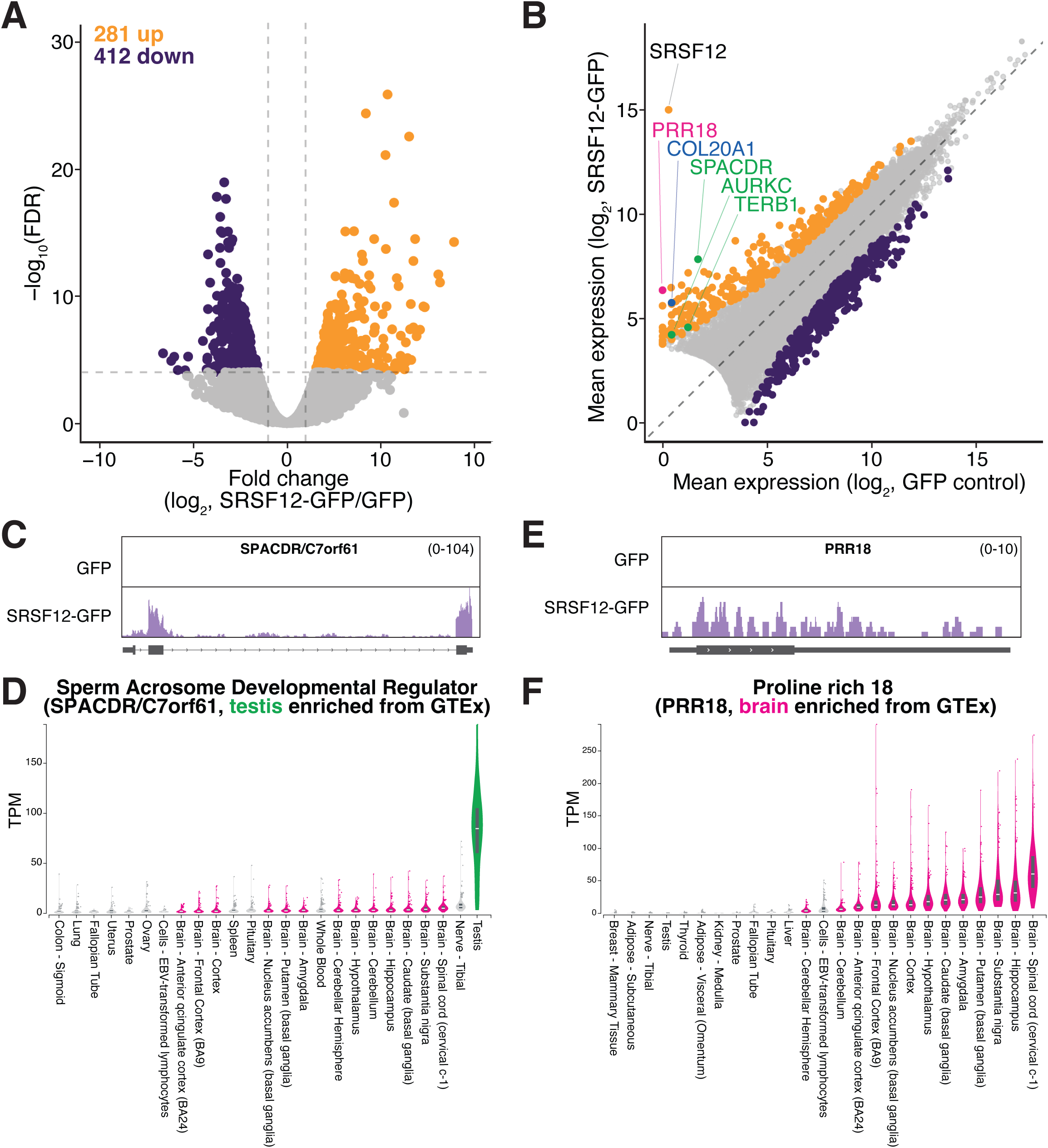
Ectopic expression of SRSF12 induces the expression of tissue specific genes. (**A**) Volcano plot of RNA-sequencing data from SRSF12-GFP vs GFP expressing cells. Orange, significantly upregulated RNAs in SRSF12-GFP; purple, significantly downregulated transcripts in SRSF12-GFP expressing cells. RNAs with false discovery rate (FDR) < 0.01 and a fold change (FC) > 2 were called as significant. n = 3 biological replicates. X and Y axis has been cropped to better highlight changes. Full plot is shown in Fig. S5A. (**B**) Scatterplot showing the mean expression of mRNAs in GFP (x-axis) and SRSF12-GFP (y-axis) expressing cells. Significance is as defined in (A). (**C**) RNA-sequencing coverage plots for SPACDR in GFP or SRSF12-GFP expressing HeLa cells. (**D**) SPACDR TPM in human tissues from (Consortium, 2013). (**E**) RNA-sequencing coverage plots for PRR18 in GFP or SRSF12-GFP expressing HeLa cells. (**F**) PRR18 TPM in human tissues from (Consortium, 2013). Top 20 most highly expressed tissues are shown.

Overall, this large-scale transcriptomic rewiring that occurs following SRSF12 expression—as marked by the inappropriate activation of typically silent gene programs in HeLa cells—may underlie the observed mitotic arrest and cell death upon SRSF12 overexpression.

## Discussion

Our results reveal that, although SRSF12 is conserved across many vertebrate species, its expression is uniquely restricted to primates where it is predominantly expressed in the testis and brain (Fig. 1). This primate-specific expression pattern raises intriguing questions about its evolutionary regulation. Whether this differential expression stems from a divergence of promoter sequences or reflects changes in the activity of upstream transcription factors between primates and other mammals remains to be determined. Here, we modeled the contributions of SRSF12 in mice, but we did not observe any clear phenotypes associated with its loss *in vivo* (Fig. S1). We speculate that this reflects the low overall expression of SRSF12 in rodents (Fig. 1C). Testing whether SRSF12 is required in primates *in vivo* will require further studies given the challenging nature of such experiments. We hypothesize that any difference between primates and rodents reflects the differences in expression levels, given the high sequence similarity between mouse and primate SRSF12 protein and the fact that mouse SRSF12 expression induced a similar phenotype in HeLa cells as human SRSF12 (Fig. S3). Alternatively, SRSF12 may be dispensable for brain or testis function in rodents through compensation by a paralog. For example, prior work suggested that SRSF10 and SRSF12 may have overlapping functions as splicing repressors (Cowper et al., 2001). Further studies are required to analyze this possible compensation from SRSF10 in SRSF12 knockout animals.

Interestingly, we found that ectopic expression of SRSF12 in HeLa cells results in a potent mitotic arrest and is lethal (Fig. 3). This lethality coincides with a large transcriptomic change, with the induction of hundreds of tissue-specific genes that are not normally expressed in HeLa cells (Fig. 6; Fig. S5). These tissue specific genes included meiosis, brain, and testis genes and may suggest a role for SRSF12 in promoting their expression in the germline and brain where SRSF12 is normally expressed in primates. Together, our work highlights the critical role of splicing regulators in organism and tissue-specific gene expression programs.

## Acknowledgments

We thank the members of the Cheeseman lab for helpful discussions; the Whitehead Genetically Engineered Models Center for mouse model generation, the Whitehead Genome Technology Core for RNA-sequencing, and the Whitehead Quantitative Proteomics Core for mass spectrometry.

## Author contributions

Conceptualization: JL and IMC; Investigation and methodology: JL performed experiments with help from EK and SH and SLC performed mouse work; Formal analysis: JL; Writing: JL and IMC wrote the manuscript; Supervision: IMC; Funding acquisition: IMC and JL

## Data and materials availability

RNA sequencing data and associated analyses were deposited in Gene Expression Omnibus (GSE303179). The mass spectrometry proteomics data have been deposited to the ProteomeXchange Consortium via the PRIDE (Perez-Riverol et al., 2025) partner repository with the dataset identifier PXD066334.

---

**Supplemental Figure 1.**
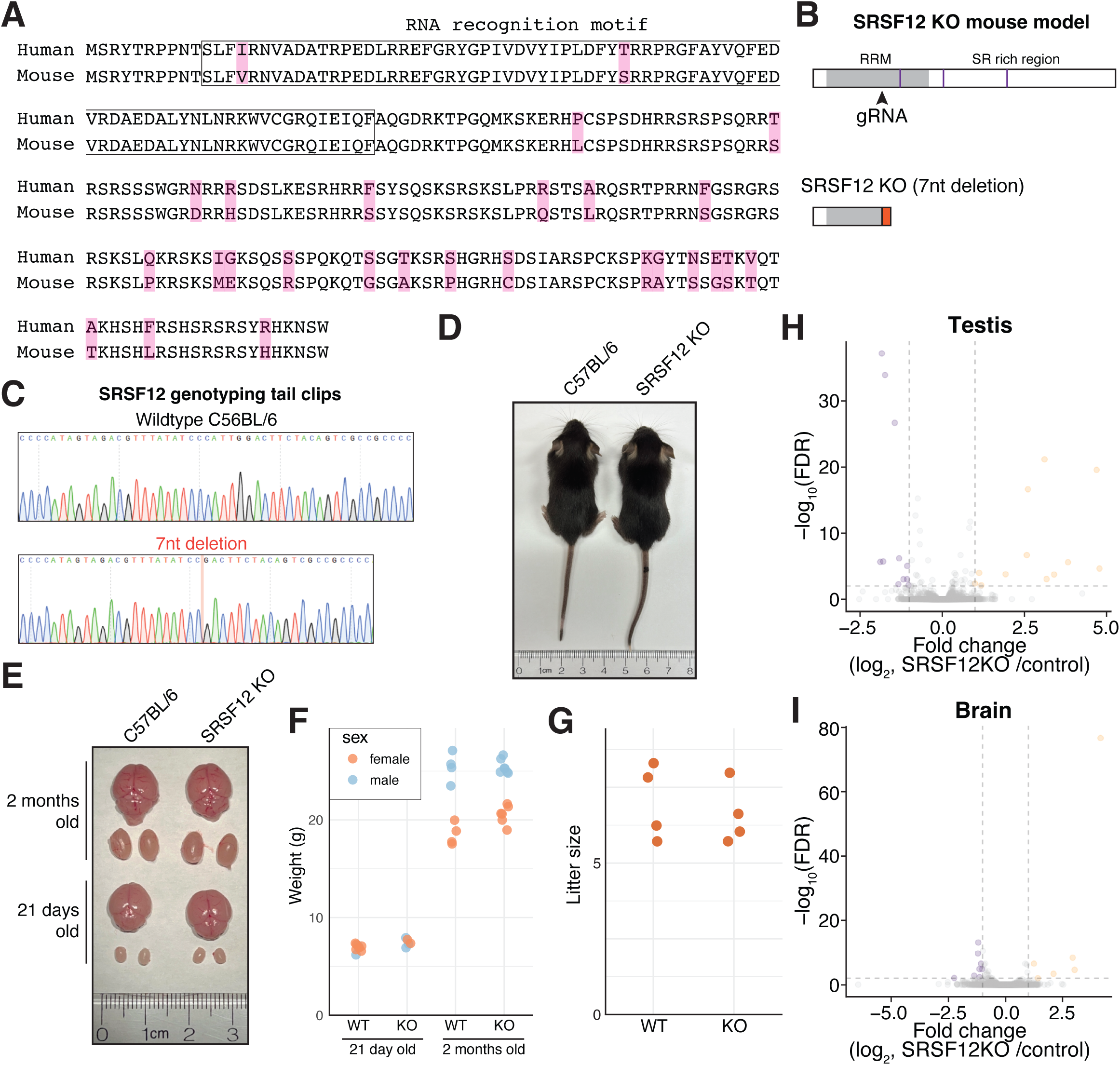
SRSF12 knockout mouse is viable with no gross defects. **(A)** Protein alignment for human and mouse SRSF12. Magenta highlight indicates differences. (**B**) Schematic of the SRSF12 mouse model. The purple lines indicate position of exon-exon junctions. The 7nt insertion generates a frameshifted SRSF12 protein starting in the middle of the RRM. (**C**) Sanger sequencing trace highlighting the 2-nt insertion or 7-nt deletion. (**D**) Representative images of wild-type and SRSF12 knockout mice at postnatal day 21. (**E**) Image of wildtype and SRSF12 KO testis and brain at the indicated age. Each dot represents a single animal (**F**) Weight of wildtype and SRSF12 knockout animals at the indicated age. (**G**) Litter sizes of wildtype and SRSF12 knockout animals. (**H**) Volcano plot for differences in RNA expression in SRSF12 knockout and control testis. n = 2 biological replicates. (**I**) Volcano plot for differences in RNA expression in SRSF12 knockout and control brain. n = 2 biological replicates.

**Supplemental Figure 2.**
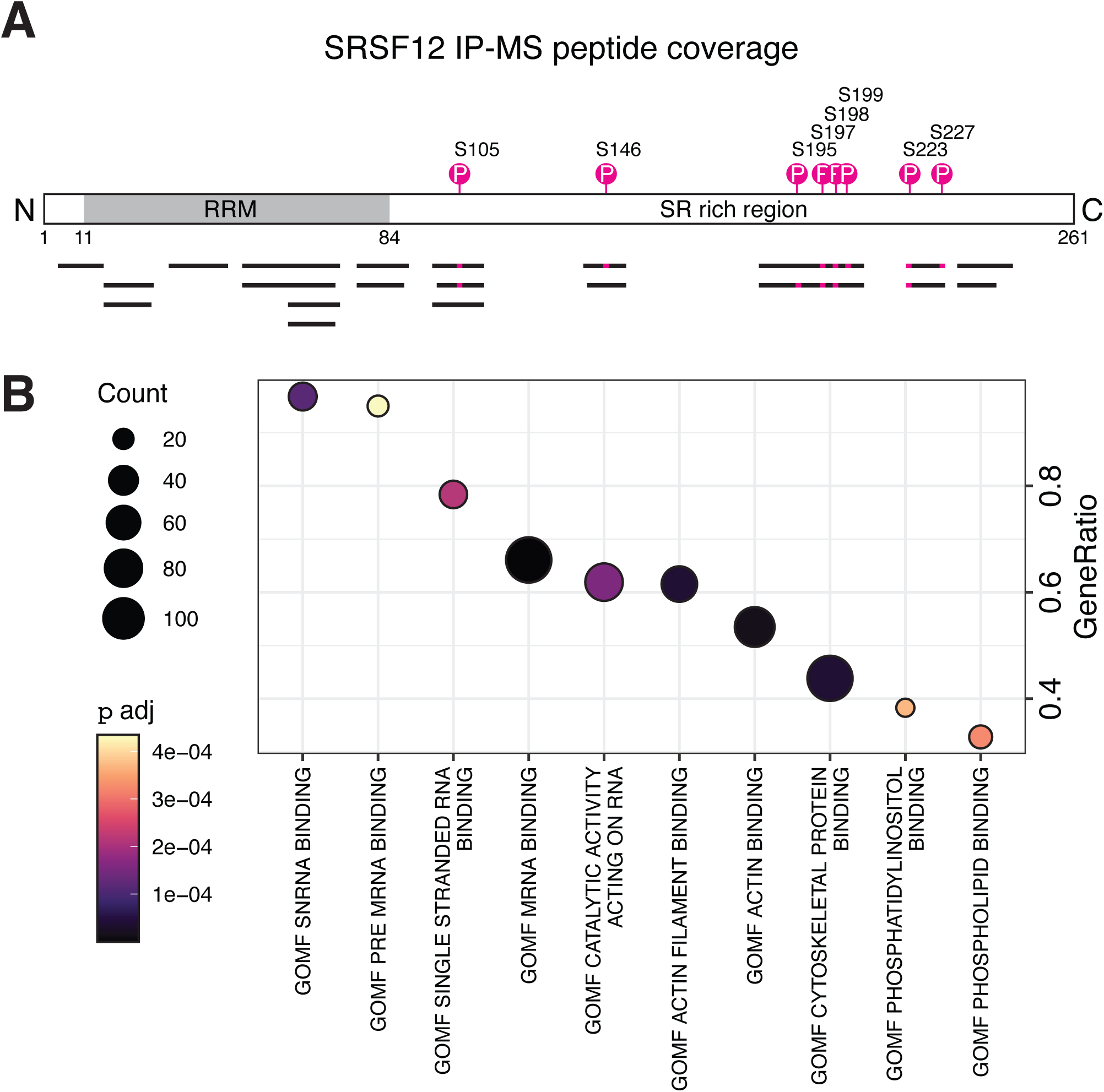
SRSF12 phosphorylation and interacting partners. **(A)** SRSF12 coverage plot from SRSF12-GFP IP-MS data. Top, a schematic representation of SRSF12. Bottom, the peptides identified in IP-MS data. Pink bars indicate phosphorylated serine residues. (**B**) GSEA was performed on SRSF12-GFP interacting proteins, and the top 10 enriched gene sets are displayed.“

**Supplemental Figure 3.**
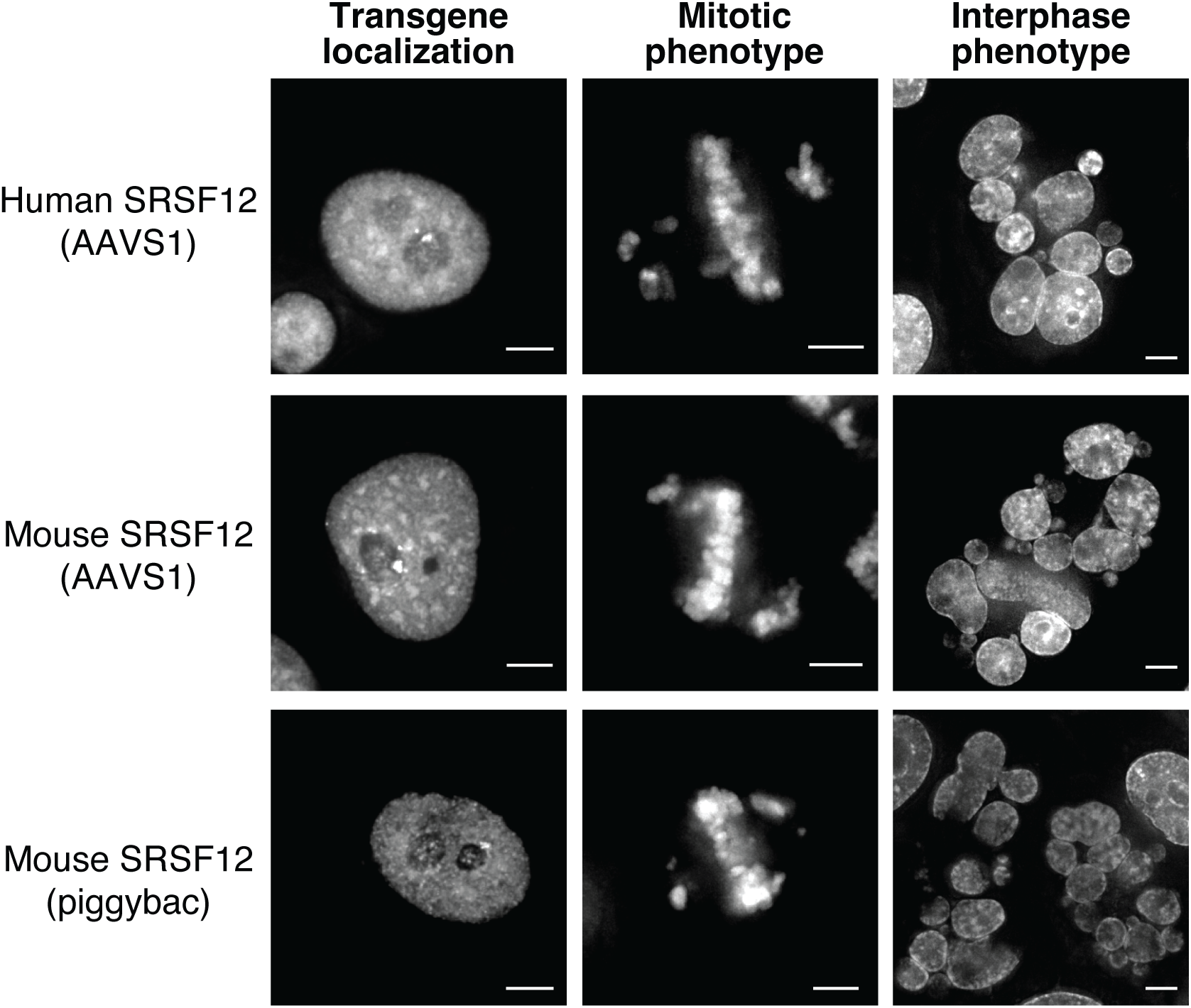
Mouse SRSF12 overexpression results in chromosome segregation errors. Left panels show the subcellular localization of SRSF12-GFP for the indicated constructs. Middle panels display representative mitotic chromosome configurations. Right panels highlight interphase phenotypes observed upon SRSF12 overexpression. Scale bar indicates 5 µm.

**Supplemental Figure 4.**
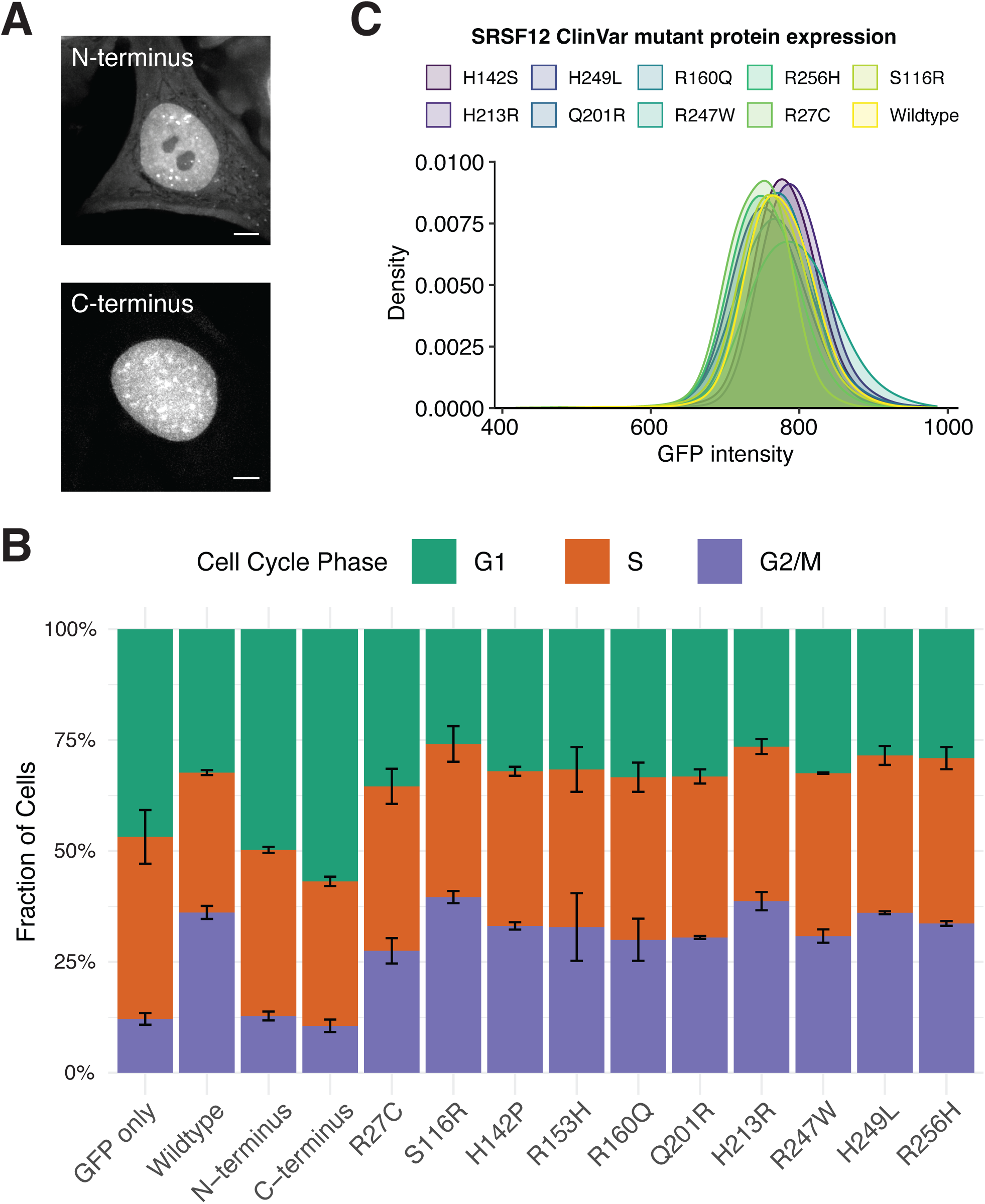
Analysis of SRSF12 ClinVar mutants. **(A)** Live cell imaging showing the localization of SRSF12 N-terminus or C-terminus tagged with GFP. Scale bar indicates 5 µm. (**B**) Stacked bar plots showing the cell cycle distribution in cells expressing each of the indicated constructs as measured by DNA-content analysis. (**C**) Density plots showing the relative levels of GFP for the different SRSF12 ClinVar mutants as measured by flow cytometry.

**Supplemental Figure 5.**
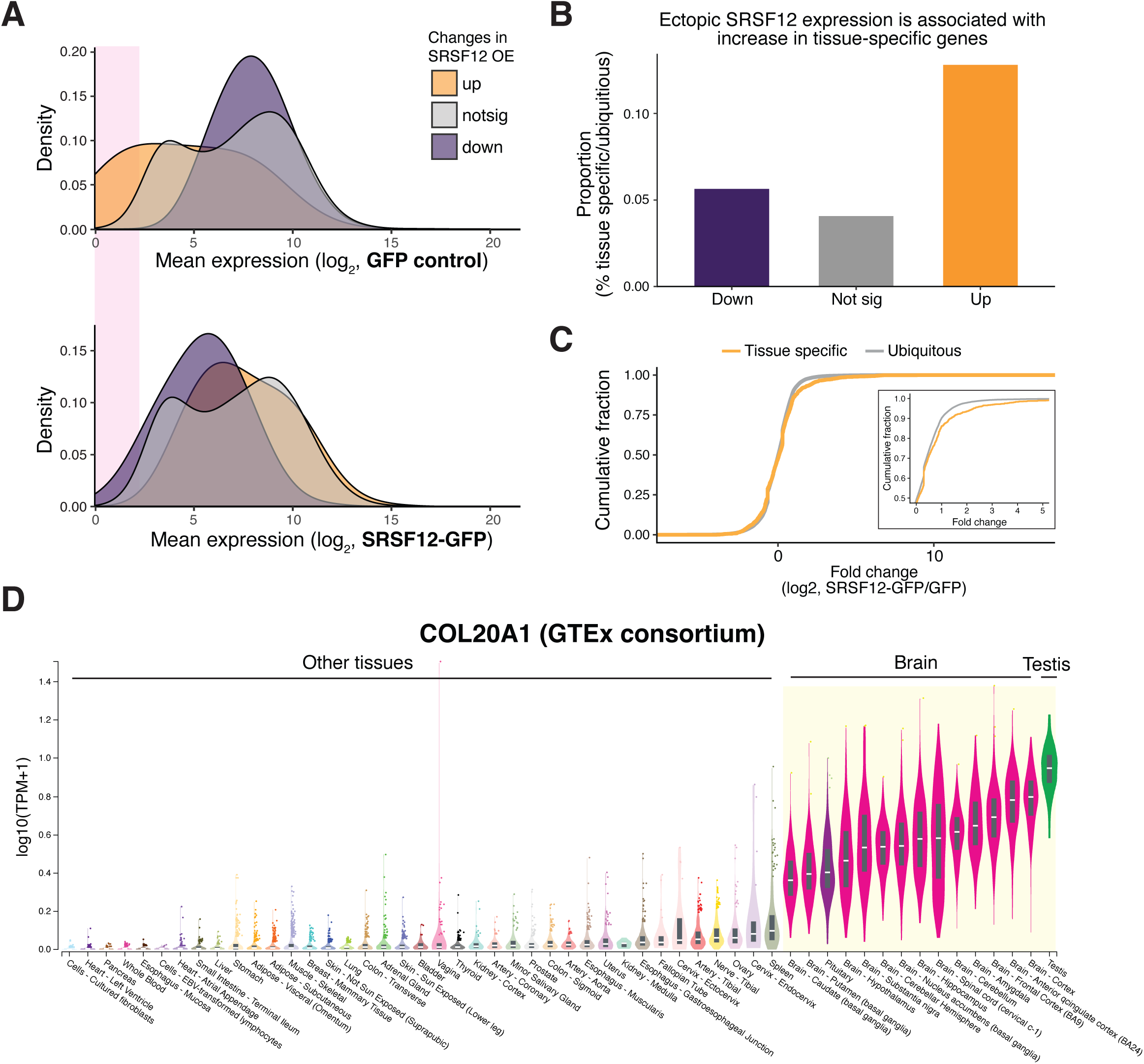
SRSF12 induced genes are lowly expressed in HeLa cells and enriched for tissue-specific genes. (**A**) Density plots showing the distribution of mean expression levels for genes altered by SRSF12-GFP expression. The top plot displays gene expression in control cells, while the bottom plot shows expression in SRSF12-GFP–expressing cells. Notably, genes upregulated by SRSF12 tend to be lowly expressed in control cells, indicating preferential activation of low-abundance transcripts. Magenta shade highlights an arbitrary region of lowly expressed genes. (**B**) Bar plots showing the relative proportion of tissue specific and ubiquitously expressed genes that are upregulated (orange), not changing (grey), or downregulated (purple) by SRSF12-GFP expression. Annotation for tissue-specific genes from the Human Protein Atlas (Uhlen et al., 2015). (**C**) Cumulative distribution function plot showing the upregulation of a subset of tissue-specific genes in cells expressing SRSF12-GFP. (**D**) COL20A1 expression (TPM+1, log_10_) from the GTEx consortium highlights the brain- and testis-enriched expression of COL20A1.

## Methods

### SRSF12 knockout mouse generation

All animals were housed in the animal facility of the Whitehead Institute and all procedures were approved by the MIT Institutional Animal Care and Use Committee (0820-020-23). C57BL/6J female mice (Jackson Laboratory) were superovulated at 4 weeks of age by intraperitoneal injection of 5 IU HyperOva (CosmoBio USA), followed 48 hours later by 5 IU human chorionic gonadotropin (hCG, ProspecBio). Oocytes were collected from oviducts approximately 16.5 hours after hCG administration and fertilized in vitro using CARD medium (CosmoBio USA) with C57BL/6J sperm capacitated in CARD FERTIUP (CosmoBio USA). Following fertilization, zygotes were cultured in EmbryoMax advanced KSOM medium (KSOM-Advanced, Millipore Sigma) at 37°C in 5% CO_₂_ until electroporation.

A synthetic single-guide RNA (sgRNA) targeting the Srsf12 locus, 5′-CGGCGACTGTAGAAGTCCAA (PAM: TGG), was obtained from Sigma-Aldrich. CRISPR/Cas9 ribonucleoprotein complexes (RNPs) consisting of 200 ng/ul Cas9 (Alt-R™ S.p. Cas9 Nuclease V3, IDT) and 100 ng/ul sgRNA were electroporated into zygotes using a CUY21EDIT electroporator (BEX Co Ltd). Zygotes were cultured overnight in KSOM-Advanced and two-cell embryos—22-29 per female—were transferred the next day to E0.5 pseudopregnant CD1 recipient females.

### Tissue culture

HeLa cells were maintained at 37°C in a humidified incubator with 5% CO_₂_, using DMEM supplemented with 10% heat-inactivated fetal bovine serum, 2 mM L-glutamine, and 100 U/mL penicillin-streptomycin.

### Molecular biology

Human and mouse SRSF12 sequences were synthesized by Twist Bioscience. With the exception of the work in Figure S2, all of the SRSF12 constructs was cloned upstream of a 3x HA tag followed by EGFP and downstream of the tetracycline inducible promoter (TRE3G). Donor plasmids contained puromycin N-acetyltransferase (puromycin resistance), reverse tetracycline-controlled transactivator, and the tetracycline inducible transgene flanked by piggyBac inverted terminal repeats. To generate stable tetracycline inducible cell lines, 1 µg donor plasmid was co-transfected with 400 ng piggyBac transposase plasmid using lipofectamine 2000. 2 days post-transfection, cells were selected with 0.45 µg/mL puromycin for at least 3 days.

We also used integrated tetracycline inducible SRSF12-MYC-TEV-EGFP at the AAVS1 site (Fig. S3) (Qian et al., 2014). To generate stable cell lines, 500 ng donor plasmid was co-transfected with 500 ng pX330-sgRNA AAVS1 using lipofectamine 2000. 2 days post-transfection, cells were selected with 0.45 µg/mL puromycin for at least 3 days.

### Analysis of previously published RNA-sequencing and ribosome profiling data

Transcript level quantification was performed using Kallisto with default settings. The TPM for each SRSF12 mRNA isoform was summed and displayed (Fig. 1D-F). For the analysis of human testis subpopulation sequencing data, we reanalyzed previously published data (Jegou et al., 2017). We reanalyzed NA-sequencing data from (Reyes et al., 2017) for aging human oocyte and eggs and (Wu et al., 2018) for GV vs MII vs ICM comparisons.

For reanalysis of NGN2-derived and HeLa cell ribosome profiling data, reads were trimmed sequentially with Cutadapt (v4.8) (Martin, 2011) to remove adapters and poly(A) tails. First, a custom 3′ adapter was trimmed with the following settings -j 20 -a ‘AGAGCACACGTCTGAACTCCAGTCACX’ -O 2 -m 14. Next, residual adapters were removed with the following settings -a ‘AGATCGGAX’ -O 1 -e 2 -m 14. Finally, poly(A) tails of ≥10 adenines were trimmed -a ‘A(Anders et al.)’ -O 3 -m 14. Trimmed reads were mapped using STAR (v2.7.1a) (Dobin et al., 2013) with the following settings -- outFilterMultimapNmax 2 --outFilterType BySJout --outSAMattributes All --outSAMtype BAM SortedByCoordinate. Reads mapping to annotated open reading frames from Gencode v25 (excluding the first 15 and last 5 codons) were quantified using HTSeq-count (v1.99.2) (Anders et al., 2015) and TPM normalized.

### RNA extraction, library prep, and sequencing

Cells expressing dox-inducible SRSF12-GFP or GFP were treated with 1 µg/mL doxycycline for 48 hours. After transgene induction, cells were rinsed once with PBS and lysed in 400 µL of TRI Reagent (Invitrogen, AM9738) before being stored at −80°C. Mouse tissues were rapidly frozen in liquid nitrogen and kept at −80°C for storage. While kept frozen on dry ice, the tissues were ground to a fine powder using a mortar and pestle to preserve cell integrity. A small portion of the pulverized frozen tissue was then suspended in 400 µL of TRI Reagent. To the TRI Reagent lysate, 120 µL of chloroform was added, followed by shaking and centrifugation at 21,000 x g for 15 minutes at 4°C to achieve phase separation. The upper aqueous phase, was carefully transferred to a new tube, mixed with an equal volume of chloroform, and centrifuged again at 21,000 x g for 1 minute at 4°C. RNA was then precipitated by adding 300 mM NaCl and 30 µg of GlycoBlue (AM9516), along with an equal volume of isopropanol, and incubating the mixture overnight at −20°C. The RNA pellet was centrifuged at 21,000 x g for 15 minutes at 4°C, washed with 70% ethanol, and resuspended in RNase-free water. RNA concentration was measured using a Nanodrop. For replicates 1 and 2 for SRSF12 overexpression and mouse tissue RNA-sequencing, ribodepletion and library preparation was done with the Watchmaker Genomics Kit. Paired end 75 x 75 sequencing was performed on an Element AVITI.

For the third replicate of SRSF12 overexpression transcriptome analysis, the total RNA was extracted using Phasemaker™ tubes according to the manufacturer’s protocol (Invitrogen, A33248). RNA was resuspended in nuclease-free water and 4 pg spike-in RNA (equal mixture of in vitro transcribed Fluc and Nluc) was added per 2 ug total RNA to capture global changes in RNA abundances. rRNA depletion and library preparation were carried out with the KAPA RNA HyperPrep Kit with RiboErase (HMR) (Roche KK8560) according to manufacturer’s instructions. Paired-end 50 x 50 sequencing was performed on an Illumina NovaSeq SP.

### RNA-sequencing analysis

Reads were mapped to the human (gencode v25, GRCh38.p7) or mouse (gencode v24, GRCm38.p6) genome using STAR with the following settings --outFilterMultimapNmax 1 --outFilterType BySJout --outSAMattributes All --outSAMtype BAM SortedByCoordinate. Exon mapping reads were quantified using htseq-count (1.99.2) with the parameters -f bam -t exon -s reverse. For SRSF12-GFP overexpressing cells --nonunique all was also included. No read cutoff was used because tissue-specific genes such as SRSF12, SPACDR, or PRR18 were not detected in control samples but were detected in SRSF12-GFP expressing cells. Differential expression analysis was performed with DESeq2 (Love et al., 2014).

Differential exon usage was analyzed using DEXSeq (Anders et al., 2012) with standard parameters. To minimize noise from lowly expressed exons, only those with a BaseMean greater than 100 were included in the analysis.

Gene set enrichment analysis (Subramanian et al., 2005) was performed using the clusterProfiler R package (v4.6.2) (Wu et al., 2021) to identify enriched Gene Ontology terms. Gene sets were obtained from the MSigDB database (v2023.1) using the msigdbr package (Liberzon et al., 2015). A gene list was ranked ordered by fold change and enrichment analysis was run with a p-value cutoff of 0.05.

We used the Human protein Atlas (Uhlen et al., 2015) to define tissue specific genes (https://www.proteinatlas.org/humanproteome/tissue/tissue+specific).

### Immunofluorescence

Dox-inducible SRSF12-GFP HeLa cells were seeded on poly-L-lysine coated coverslips. One day after seeding, the transgene was induced for 48 hours. The cells were pre-extracted using PBS + 0.25% Triton X-100 for 2 minutes at room temperature and fixed in PBS + 4% formaldehyde + 0.25% Triton X-100 at room temperature for 15 minutes. Following fixation, cells were washed 3 times with PBS + 0.1% Triton X-100 and blocked in AbDil for 30 minutes-1 hour. Following blocking, cells were stained at 4°C for 16 hours with α-SRSF1 antibody (1:500, Proteintech, 12929-2-AP) diluted in AbDil. Cells were washed with PBS + 0.1% Triton X-100, 3 times then incubated in α-GFP nanobody conjugated to AlexaFluor488 (1:1000, Proteintech, gb2AF488) and Donkey α-Rabbit IgG (H+L) AlexaFluor647 secondary antibody (1:300, Jackson ImmunoResearch Laboratories) diluted in AbDil for 1 hour at room temperature. After secondary, cells were stained in hoescht for 15 minutes at room temp, then washed 3 times with PBS + 0.1% Triton X-100 before mounting in PPDM (0.5% p-phenylenediamine and 20 mM Tris-Cl, pH 8.8, in 90% glycerol) and sealed with nail polish.

### Microscopy

The GFP transgene was induced by addition of 1 µg/mL dox for 24 hours prior to imaging. Cells on glass bottom plates were incubated in 0.1 μg/mL Hoescht for >30 minutes. All images were taken on the Deltavision Ultra (Cytiva) system using a 60x/1.42NA objective. 8 μm images were taken with z-sections of 0.2 μm. All images were deconvolved and max projected. Since we did not quantitatively compare intensities between images and sought to evaluate subcellular localization generally, the images were not scaled equivalently.

### Cell cycle analysis using flow cytometry

Cells expressing the SRSF12 transgene were seeded in 6-well plates and induced for 48 hours as previously described. After induction, culture media was transferred to a 15 mL conical tube. Adherent cells were gently detached by treatment with PBS containing 5 mM EDTA and then combined with the harvested media. Cells were collected by centrifugation at 500 × g, washed once with 1X PBS, and resuspended in 0.5 mL PBS. Ice-cold ethanol was added dropwise to the suspension while gently mixing, and cells were fixed at 4 °C for at least 30 minutes.

Following fixation, cells were washed once with PBS containing 0.3% BSA (w/v), then again with PBS containing 3% BSA (w/v) and 0.1% Triton X-100 (v/v). Blocking was performed by incubating cells at room temperature for 30 minutes in antibody dilution buffer (AbDil: 20 mM Tris-HCl, 150 mM NaCl, 0.1% Triton X-100, 3% BSA, 0.1% sodium azide, pH 7.5). Cells were then incubated overnight at 4 °C with end-to-end rotation in a 1:1,000 dilution of anti-phospho-histone H3 Ser10 antibody (Abcam, ab5176) prepared in AbDil. The next day, cells were washed with AbDil three times then incubated for 1 hour at room temperature with 1:1,000 α-GFP nanobody conjugated to AlexaFluor488 (Proteintech, gb2AF488) and 1:300 goat anti-rabbit Cy5 (Jackson ImmunoResearch Laboratories), both diluted in AbDil. After staining, cells were washed three times with AbDil, then incubated for 30 minutes at room temperature in AbDil supplemented with 20 μg/mL Hoechst.

Cells were filtered through a cell strainer before acquisition on a BD FACSymphony A1 Cell Analyzer (BD Biosciences). Flow cytometry data were processed using FlowJo software. Quantification was performed after gating for live, single cells. The cell cycle tool with the Watson model was used to quantify the relative proportion of G1, S, G2/M cells based on DNA content. To quantify mitotic cells, we gated for phosphorylated histone H3 Serine 10 positive cells.

### Competitive growth assay

Equal amounts of control dox-inducible mCherry cells were mixed with dox-inducible GFP or SRSF12-GFP cells. The population of GFP:mCherry cells were monitored every few days using the BD FACSymphony A1 Cell Analyzer (BD Biosciences). Flow cytometry data was analyzed using FlowJo (v10.7.1).

### Immunoprecipitation

Cells were detached by incubating them in 1X PBS with 5 mM EDTA for 10 minutes at 37 °C. After collection, cells were pelleted by centrifugation at 200 x g, washed once with 1X PBS, followed by a second wash with lysis buffer (25 mM HEPES pH 8.0, 2 mM MgCl_₂_, 0.1 mM EDTA pH 8.0, 0.5 mM EGTA pH 8.0, 300 mM KCl, 10% glycerol), with centrifugation between washes. The final pellet was resuspended in an equal volume of 1X lysis buffer, snap frozen in liquid nitrogen, and stored at −80 °C.

To each frozen pellet, an equal volume of polysome lysis buffer (20 mM HEPES pH 7.5, 100 mM KCl, 5 mM MgCl_₂_, 1% Triton X-100) supplemented with 1% CHAPS, 1× cOmplete EDTA-free protease inhibitor cocktail (Roche), and 0.02 U/μl SUPERase•In (Invitrogen) was added prior to thawing at 37 °C. Cell disruption was performed via sonication using a Bioruptor (Diagenode) at high amplitude for 10 cycles (30 seconds on, 1 minute off). Lysates were clarified by centrifugation at 21,000 x g for 30 minutes at 4 °C.

The supernatant was incubated with Protein A magnetic beads (Bio-Rad) pre-conjugated to rabbit anti-GFP antibody (Cheeseman and Desai, 2005) for 2 hours at 4 °C on a rotating wheel. Beads were then washed four times with polysome lysis buffer supplemented with 1 mM DTT, 10 µg/ml leupeptin, pepstatin, and chymostatin, and 0.02 U/μl SUPERaseIn. Bound proteins were eluted three times using 2X bead volumes of 100 mM glycine pH 2.6 for 5 minutes each at 4 °C. Eluates were pooled, neutralized with Tris pH 8.5 to a final concentration of 200 mM, and proteins were precipitated overnight at 4 °C by adding trichloroacetic acid to 1/5 of the final volume. Precipitated proteins were pelleted by centrifugation (21,000 x g, 30 minutes), washed three times with 1 mL ice-cold acetone, and dried using a speedvac for 5 minutes at 30 °C.

### Mass spectrometry sample preparation

Protein pellets were resuspended in 5% SDS, 50 mM TEAB pH 8.5, and 20 mM DTT, then heated at 95 °C for 10 minutes. After cooling to room temperature, iodoacetamide was added to a final concentration of 40 mM and incubated for 30 minutes in the dark. The reaction was quenched by adding 2.5% phosphoric acid (v/v), and 6 volumes of S-Trap binding buffer (90% methanol, 100 mM TEAB, pH 7.55) were added.

Samples were loaded onto S-Trap microcolumns (Protifi) by centrifugation at 4000 x g for 1 minute. The flow-through was reapplied, and columns were washed four times with S-Trap binding buffer. On-column digestion was carried out by adding 1 µg trypsin in 50 mM TEAB pH 8.5 and incubating at 37 °C overnight in a humidified chamber.

Tryptic peptides were sequentially eluted with 40 µL of 50 mM TEAB, 40 µL of 0.2% formic acid, and 35 µL of 50% acetonitrile/0.2% formic acid. Pooled eluates were flash frozen and lyophilized for 4 hours. Peptide concentration was measured using the Pierce Fluorometric Peptide Assay Kit according to the manufacturer’s instructions. Samples were reconstituted in 0.1% formic acid to a final concentration of 250 ng/μL, and 250 ng of peptide was injected into an Exploris 480 Orbitrap mass spectrometer for analysis.

### Mass spectrometry and analysis

Raw mass spectrometry data were analyzed using Proteome Discoverer version 2.4 (Thermo Fisher Scientific). Protein and peptide identification was performed with the Sequest HT algorithm (Eng et al., 1994), using a human reference proteome (UP000005640) supplemented with the GFP sequence. Searches allowed for tryptic digestion with up to two missed cleavages. Precursor ion mass tolerance was set to 10 ppm, and fragment ion tolerance was set at 0.02 Da. The search parameters included dynamic modifications for phosphorylation (+79.966 Da on serine, threonine, and tyrosine), methionine oxidation (+15.995 Da), and N-terminal acetylation (+42.011 Da), along with a static modification for carbamidomethylation on cysteines (+57.021 Da). To control for false discoveries, peptide-spectrum matches were filtered using Percolator, applying a threshold of <1% false discovery rate.

For the analysis of SRSF12 interacting partners, we only included proteins with 3 or more unique peptides. We excluded mitochondrial proteins found in the immunoprecipitations which represented in vitro reassociation, given the strict nuclear localization of SRSF12.

